# snakePipes enable flexible, scalable and integrative epigenomic analysis

**DOI:** 10.1101/407312

**Authors:** Vivek Bhardwaj, Steffen Heyne, Katarzyna Sikora, Leily Rabbani, Michael Rauer, Fabian Kilpert, Andreas S Richter, Devon P Ryan, Thomas Manke

## Abstract

The scale and diversity of epigenomics data has been rapidly increasing and ever more studies now present analyses of data from multiple epigenomic techniques. Performing such integrative analysis is time-consuming, especially for exploratory research, since there are currently no pipelines available that allow fast processing of datasets from multiple epigenomic assays while also allow for flexibility in running or upgrading the workflows. Here we present a solution to this problem: snakePipes, which can process and perform downstream analysis of data from all common epigenomic techniques (ChIP-seq, RNA-seq, Bisulfite-seq, ATAC-seq, Hi-C and single-cell RNA-seq) in a single package. We demonstrate how snakePipes can simplify integrative analysis by reproducing and extending the results from a recently published large-scale epigenomics study with a few simple commands. snakePipes are available under an open-source license at https://github.com/maxplanck-ie/snakepipes.

## Main

Epigenomics is a fast growing field, and due to the consistent fall in the price of sequencing, increase in multiplexing abilities, and multiple innovations in laboratory protocols, it has become increasingly convenient to perform multiple epigenomic assays within a project. However, a major bottleneck on the way to process and analyse this data in a reproducible way, particularly for novice analysts, is the availability of analysis pipelines. Next-generation sequencing (NGS) analysis pipelines are composed of a series of data processing steps, employ standardised processing parameters, and are usually scalable to large number of samples ^1^. Due to such properties, most pipelines are currently developed and deployed for settings where standardized, large scale analysis is required. Examples are RNA-seq variant-calling pipelines deployed in clinical settings ^2^, or processing pipelines developed for large-scale consortia ^3,4^.

However, in a typical basic science research setting, researchers also seek to modify parameters, update tool versions or extend the workflows, while maintaining their scalability and ease-of-use. Conventional NGS pipelines, although scalable, do not allow for this flexibility. Options for exploratory and downstream analysis have been limited, resulting in various expert users developing their own custom pipelines suited to their needs. Computational frameworks such as Galaxy ^5^ and Nextflow ^6^ exist, but they still demand novice users to be trained and implement their workflows themselves from scratch. This leads to a conundrum, how can we provide a set of workflows following best-practices that are easy to install and run for the novice users, while still providing the flexibility of extending and upgrading the workflows to the expert users?

We developed snakePipes to address such requirements. snakePipes provide flexible processing as well as downstream analysis of data from the most common assays used in epigenomic studies: ChIP-seq, RNA-seq, whole-genome bisulfite-seq (WGBS), ATAC-seq, Hi-C and single-cell RNA-seq in a single package (Fig. 1b, Implementation Details). It employs snakemake ^7^ as a workflow language, which benefits from easy readability of the code, widespread adoption, and offers scalability using most cluster and cloud computing platforms. snakePipes also makes use of conda environments and the bioconda platform ^8^, which allow hassle-free installation and upgrade of tools (Fig. 1a, Implementation Details). Conda environments allow execution of tools avoiding dependency conflicts, and do not require root permissions to run. Due to a modular architecture, various tools are shared between workflows, which simplifies data integration since data from multiple technologies are processed using identical tool versions and genome annotations. The genome annotations and indices are shared by all workflows, and can also be generated directly via snakePipes, facilitating easy setup as well as integrative analysis. Finally, workflows in snakePipes employ extensive quality-checks and also produce reports using multiQC ^9^ and R, that inform the user of processing and analysis results.

**Figure 1.**
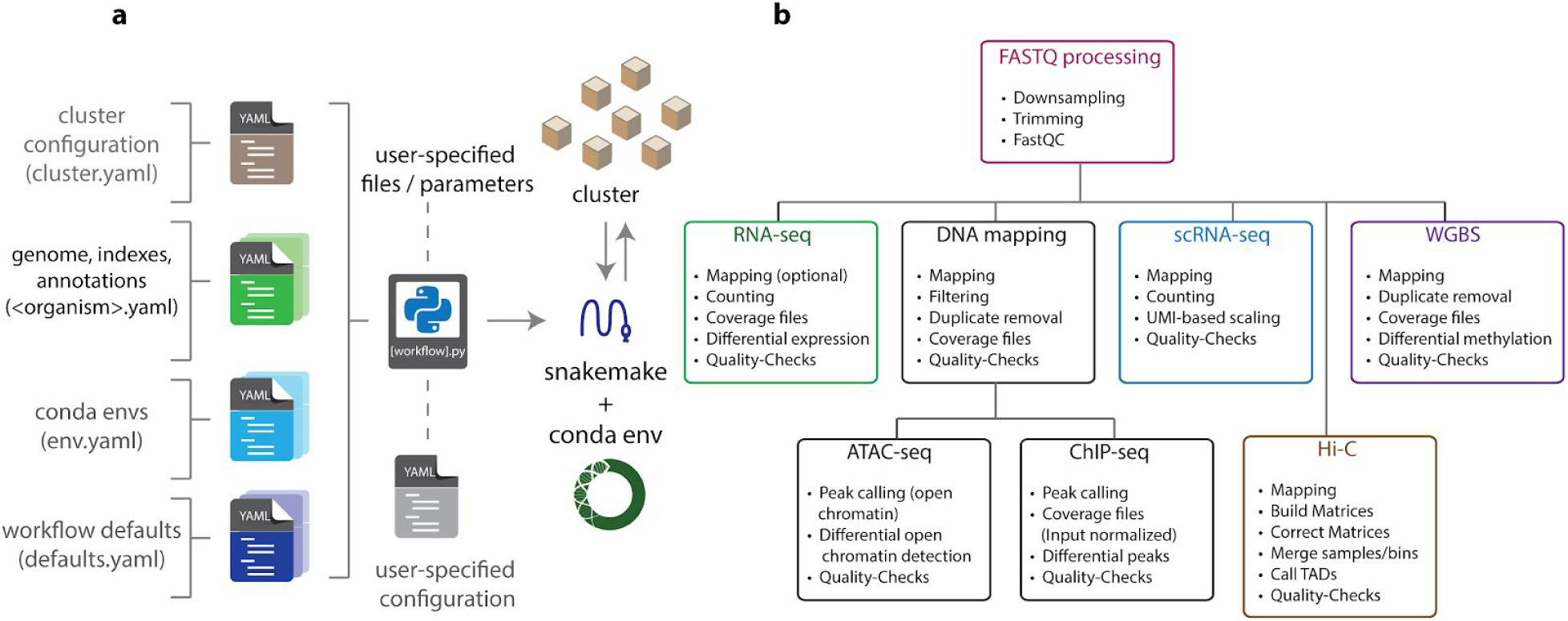
General architecture and available workflows in snakePipes. a. Configuration. All configurable parameters for snakepipes are defined as YAML files. Organism-specific YAML files need to be configured once during setup, while the configuration of cluster, conda envs and workflow defaults is optional. After installation, the location of these files can be revealed by the snakePipes info command. Workflows are all executed using their command line wrappers. During execution user parameters and/or a user-provided YAML file can be used to override the workflow defaults and add flexibility to the processing/analysis. b. Execution. FASTQ files provided by the user first goes through FASTQ processing, followed by one of the workflows. Outputs from DNA mapping workflow can be further used as input for the ChIP-seq and ATAC-seq workflows. Only general processing steps are listed and each workflow includes workflow-specific quality-checks.

Apart from conventional processing steps such as mapping, counting and peak calling, workflows in snakePipes also include various downstream analysis. All workflows (except scRNA-seq workflow) optionally accept a sample information (tab-separated) file that can be used to define groups of sample. This allows comparative analysis such as differential gene or transcript expression analysis for the RNA-seq workflow, differential peak calling for ChIP-Seq workflow, differential open chromatin detection for ATAC-seq workflow, and detection of differentially methylated regions (either de-novo or on user-specified regions) for WGBS workflow. The HiC workflow uses sample information to merge groups and can perform TAD calling with parameters adapted to the resolution of produced matrix (using HiCExplorer ^10^). This preliminary analysis, combined with visualization-ready bed and bigwig files, allows users to quickly interpret their data.

Most workflows in snakePipes also allow processing and downstream analysis of data in an allele-specific manner. DNA-mapping and RNA-seq workflows utilize SNPSplit ^11^ to map data to a single or dual-hybrid genome, for samples coming from inbred mouse strains or from *Drosophila* SNP lines using the “allelic-mapping” mode. Single/dual hybrid genome indices can be provided externally, or can be created on the fly by simply providing a VCF file ^12^ and the name of desired strains. The workflows also generate allele-specific coverage files, QC reports, and performs allele-specific differential analysis using appropriate statistical design, without requiring any additional user intervention.

In order to demonstrate how snakePipes can simplify analysis and interpretation of data from multiple epigenomic assays, we downloaded and processed data from from a recently published study that investigated the role of Smchd1 protein on the mammalian X-chromosome ^13^. The knock-out of Smchd1 in mouse neural progenitor cells (NPCs) affects the organization of inactive X-chromosome and leads to a loss of H3K27me3 domains along with a gain of H3K4me3 on various genes on the inactive X-chromosome. This results in de-repression of genes, seen as up-regulation in RNA-seq data. We observed the same effects upon re-analysis of the ChIP-Seq, RNA-seq and Hi-C data from the study, directly from the output of snakePipes, without further downstream analysis (Fig 2, Fig S1a-b). Further, we integrated this information with ATAC-seq data available online ^14^ to discover that the genes de-repressed upon Smchd1 knock-out display significantly higher open chromatin signal at their promoters in the wild-type NPCs, compared to down-regulated or unchanged, lowly expressed genes (Fig. S1c). Whole-genome bisulfite data obtained from another study (GSE101090) suggested that under wild-type conditions, promoters of the de-repressed genes show methylation levels slightly higher than the downregulated genes but significantly lower than unchanged, lowly expressed genes (Fig. S1d), corroborating previous ^15^ and recent ^16^ links between promoter CpG methylation and gene repression.

**Figure 2.**
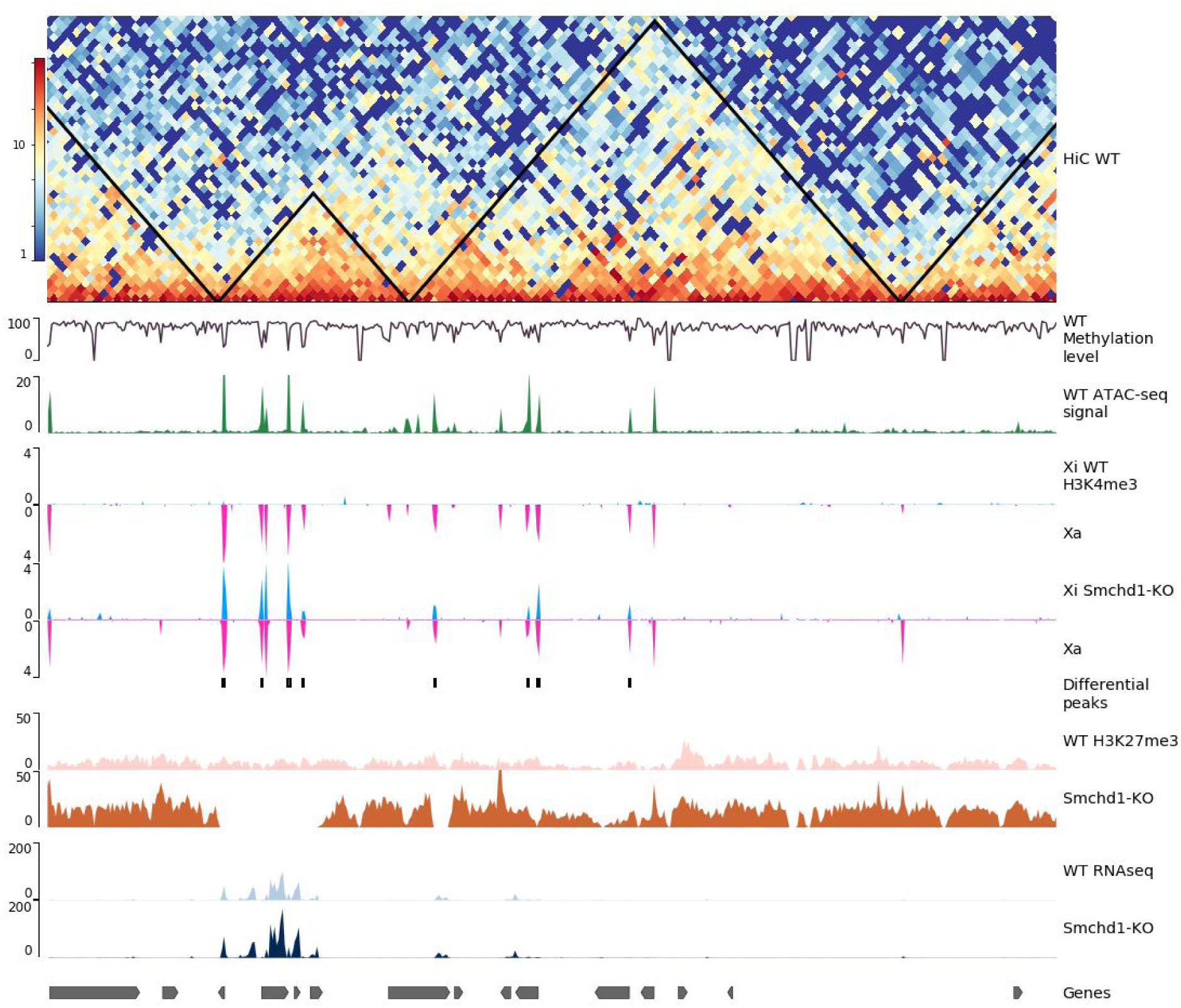
Outputs from snakePipes workflows quickly provide meaningful biological interpretation. Genome tracks plotted using pyGenomeTracks^10^. **Top track:** output of HiC workflow with TADs (black triangles) on wild-type (WT) NPCs. **Tracks 2:** Methylation signal reported from the whole-genome bisulfite-seq (WGBS) workflow on wild-type NPCs. Gene promoters show demethylation. **Track 3**: Open chromatin signal from ATAC-seq workflow on wild-type NPCs. **Track 4-7:** output of allele-specific ChIP-Seq workflow, Inactive X chromosome (Xi) tracks are in Blue while the active X chromosome (Xa) is shown in Pink. The knock-out (KO) of Smchd1 shows increase in H3K4me3 on Xi. **Track 8:** Allele-specific differential peaks detected by the ChIP-Seq workflow in KO, over WT. **Track 9-10:** H3K27me3 signal from the ChIP-Seq workflow. KO shows loss of H3K27me3 domains on certain genes. **Track 11-12:** RPKM signal from RNA-seq workflow. Genes which loose H3K27me3 show up-regulation in KO. **Bottom track:** Genes.

Our results demonstrate how a multi-assay epigenomic analysis toolkit like snakePipes could simplify data processing, reproduce previously published results, and allow new biological interpretations with minimal effort. snakePipes are under active development, open source and available via conda: ‘***conda install -c mpi-ie -c bioconda -c conda-forge snakePipes***’. Source code is available at https://github.com/maxplanck-ie/snakepipes and the documentation is hosted at https://snakepipes.readthedocs.io/en/latest/.

## Implementation Details

### General architecture of snakePipes

The general architecture of snakePipes is summarized in Fig. 1a. snakePipes utilizes conda and bioconda for setup and execution of workflows. All information required for workflow execution is stored in easy-to-edit YAML (Yet Another Markup Language) files. The **cluster.yaml** file defines the command used to execute each process on a cluster (assuming HPC cluster infrastructure like Slurm or SGE has been set up). For each organism of interest, the **organism>.yaml** file describes the location of genome fasta, (mapping) indices and annotations. This allows running different workflows on exactly the same version of genome and annotations. The conda **env.yaml** file is used to specify version number of required tools for each workflow. The required tools are then fetched and setup automatically via the conda and bioconda repositories either during workflow setup, or execution. The **defaults.yaml** file specifies reasonable default parameter for each workflow according to the best practices for the most common sequencing protocols. These files are used by the command line wrappers, that use snakemake to execute the workflows on the cluster. In the absence of cluster setup, workflows can also be executed locally. Each step of a workflow is defined as a snakemake “rule”, which is executed in it’s own virtual environment, avoiding conflicts between tools. All log files, including user-supplied commands are written in the working directory, along with (optionally) a graph of executed steps. This allows users to easily reproduce and communicate their analysis in the future.

#### Running and testing the workflows

Comprehensive documentation for snakePipes can be found online: https://snakepipes.readthedocs.io/ and the test datasets are available on zenodo: https://zenodo.org/record/1346303. snakePipes also provide a ‘***createIndices***’ workflow that creates genome indices and annotations (contents of <organism>.yaml) from a user-specified genome fasta file or URL.

## Workflows in snakePipes

snakePipes provide DNA-mapping, ChIP-seq, ATAC-seq, RNA-seq, whole-genome bisulfite-seq (WGBS), HiC and single-cell RNA-seq workflows. All workflows make use of the common fastq downsampling (via seqtk (https://github.com/lh3/seqtk)) and trimming (via cutadapt ^17^ and Trim Galore! (https://www.bioinformatics.babraham.ac.uk/projects/trim_galore/)) module. All workflows also produce an interactive report using multiQC ^9^ that summarizes outputs from multiple workflow steps and samples.

In the **DNA-mapping** workflow, the fastq files are aligned to the genome via Bowtie2 ^18^ and filtering can be performed via samtools ^19^ using user-provided parameters. Various quality-checks are performed via SamBamba ^20^, Picard ^21^, deepTools ^22^ and (optionally) qualimap ^23^. Coverage files (bigwigs) are generated via deepTools. The output of DNA-mapping workflow can then be used for ChIP-Seq or ATAC-seq workflows. DNA-mapping workflow handles both single and paired-end files and could also be used for whole-genome alignments.

The **ChIP-seq** workflow takes information about the samples (corresponding input controls, expected broad/sharp mark) using a yaml file, and performs ChIP-specific quality-checks via deepTools. It also generates input-normalized bigwig files and performs peak calling for both sharp (via MACS2 ^24^) and broad (via histoneHMM ^25^) marks. The **ATAC-seq** workflow takes paired-end DNA mapping output and performs quality-checks and filtering useful for ATAC-seq samples. It then performs detection of open chromatin using MACS2. Both ChIP-Seq and ATAC-Seq workflows can perform detection of differential peaks or differential open chromatin regions between groups of samples using CSAW ^26^, if a sample sheet is provided.

The **RNA-seq** workflow can be run in “alignment” or “alignment*-free*” mode. In the alignment mode, fastq files are aligned to the genome via user-selected aligner (STAR ^27^ or HISAT2 ^28^) and high-quality primary alignments are counted via featureCounts ^29^. In the alignment-free mode, the transcripts are directly quantified via Salmon ^30^. Transcripts can be filtered for various features before quantification. Additionally, the “*deepTools_qc*” mode can be added, which performs various quality-checks via deepTools and produces normal and depth-normalized (RPKM) coverage files. Differential gene and transcript expression analysis could then be performed using DESeq2 ^31^, wasabi (https://github.com/COMBINE-lab/wasabi) and Sleuth ^32^.

The **scRNA-seq** workflow performs mapping and counting of data obtained from the CEL-Seq2 protocol ^33^. Fastq files are first preprocessed by moving cell barcodes and unique molecular indices (UMIs) to read headers and then mapped using STAR (Dobin et al.). Quantification is then performed per-cell by accounting for UMIs and the resulting counts corrected for Poisson sampling. A variety of quality control steps are also taken, such as computing a heatmap per well-plate of obtained transcript counts and the correlation between reads and UMIs. After the workflow is finished the resulting counts files are ready for custom downstream analysis (e.g., clustering or differential expression).

The **Hi-C** workflow performs read mapping of paired-end HiC data using BWA ^34^. It then uses HiCExplorer ^10^ to build and correct the Hi-C matrices using the iterative correction (ICE) method ^35^. Matrices can be built at a user-specified resolution or at restriction fragment resolution by simply specifying the name of the restriction enzyme. The corrected HiC matrices can then be used for detection and visualization of topologically associated domains (TADs) ^36^. Quality reports are produced using HiCExplorer and are then summarized by MultiQC for comparison of samples.

The **WGBS** (whole-genome bisulfite-seq) workflow performs mapping of paired-end WGBS-seq data on a bisulfite converted genome using bwa-meth ^37^. To help assess the quality of the experiment, several metrics, including bisulfite conversion rate and coverage of random CpGs, are collected in a report. Counting of reads supporting methylated and unmethylated cytosines is performed with MethylDackel (https://github.com/dpryan79/MethylDackel). De novo discovery of differentially methylated regions (DMRs) can also be performed using Metilene ^38^ if a sample sheet is provided.

## Methods

### Processing of Online data

HiC, ChIP-Seq, and RNA-seq data for Smchd KO and wild-type Neural Progenitor Cells (NPCs) was downloaded from GSE99991. ATAC-Seq data for wild-type NPCs was downloaded from GSE71156 and WGBS data for wild-type NPCs was downloaded from GSE101090. All data was processed with snakePipes (version 1.0.0alpha5) on mouse genome GRCm38 (mm10).

The parameters specified for processing the data are as follows:

#### ATAC-Seq

**DNA-mapping** performed on mouse genome with parameters: *‘-m allelic-mapping -j 30 --gcbias --mapq 5 --dedup --fastqc --trim --properpairs*

**ATAC-seq** performed on DNA-mapping output with parameters: *‘--bw-binsize 10‘.* By default, fragments longer than 150 nt are removed from peak calling *‘--atac-fragment-cutoff 150‘*.

#### ChIP-Seq

**DNA-mapping** performed on a dual hybrid (129S1/CAST) mouse genome with parameters: *‘-m allelic-mapping --trim --fastqc --bw-binsize 10 --plotFormat pdf --dedup --mapq 10 --SNPfile <snp_positions.txt> --Nmasked_index <bowtie2_index.bt2> ‘*

**ChIP-Seq** performed on DNA-mapping output with parameters: ‘--bw-binsize 10‘ with

*chip_sampleInfo.yaml* file which specified the corresponding input controls and peak-type. H3K4me3 samples were specified as ‘*sharp‘* while the H3K27me3 samples were described as ‘*broad‘*

#### RNA-Seq

**RNA-seq** workflow was run with parameters: ‘--fastqc --trim -m alignment, deepTools_qc --DE sampleinfo.tsv’ where sampleinfo.tsv file defined groups with replicates (5 control and 5 knock-out samples). One of the knock-out samples was removed after inspecting PCA output (Fig. S1A) and workflow was re-run to obtain differentially expressed genes (Fig. S1B).

#### Hi-C

**Hi-C** workflow was run with parameters: ‘*--merge_samples --sampleInfo sampleinfo.tsv --distVsCount --bin_size 10000 --trim --fastqc*‘ where sampleinfo.tsv was used to define the two Hi-C replicates.

#### WGBS

**WGBS** workflow was run with all default parameters.

## Downstream analysis

Plotting of ATAC-seq signal on up-regulated, down-regulated and 500 lowly expressed, un-affected genes was performed by subsetting gene sets from DESeq2 output of snakePipes RNA-seq module, followed by deepTools computeMatrix with options: *-a 5000 -b 5000*, and plotHeatmap with option *--plotType se*. For WGBS data, bedgraph output of CpG methylation signal from WGBS pipeline was converted to bigWig using *UCSCtools bedGraphToBigWig* and plotting using computeMatrix (--binsize 100) and plotHeatmap.

## Code availability

snakePipes is open-source and freely available on GitHub: https://github.com/maxplanck-ie/snakepipes

## Acknowledgements

Authors acknowledge the contribution of Gina Renschler and Jana Böhm on testing of workflows. We also thank Chen-Yu Wang for useful comments on our preprint. TM acknowledges funding from the German Science Foundation (CRC992 “Medical Epigenetics”).

## Availability of data and materials

Online datasets re-analysed during this study are available in GEO with accession numbers: GSE99991, GSE71156 and GSE101090

## Authors’ contributions

VB developed the allele-specific and HiC workflows and contributed to DNA-mapping, ChIP-seq, ATAC-seq and RNA-seq workflows and documentation. SH developed the scRNA-seq workflow and documentation and contributed to DNA-mapping, ChIP-seq and RNA-seq workflows. DPR improved the wrapper design, integrated installation and conda support, contributed to the documentation and bug fixes to various workflows. KS developed the WGBS workflow and documentation. LR contributed to HiC workflow and documentation. MR developed ATAC-seq workflow. FK contributed to RNA-seq workflow. AR contributed to DNA-mapping and ChIP-seq workflow. FK, SH and AR contributed to the general design of snakePipes and wrote the early version of the wrappers. VB performed the analysis and wrote the manuscript with input from all authors. TM conceived the project and supervised the development of snakePipes.

---

**Figure S1.**
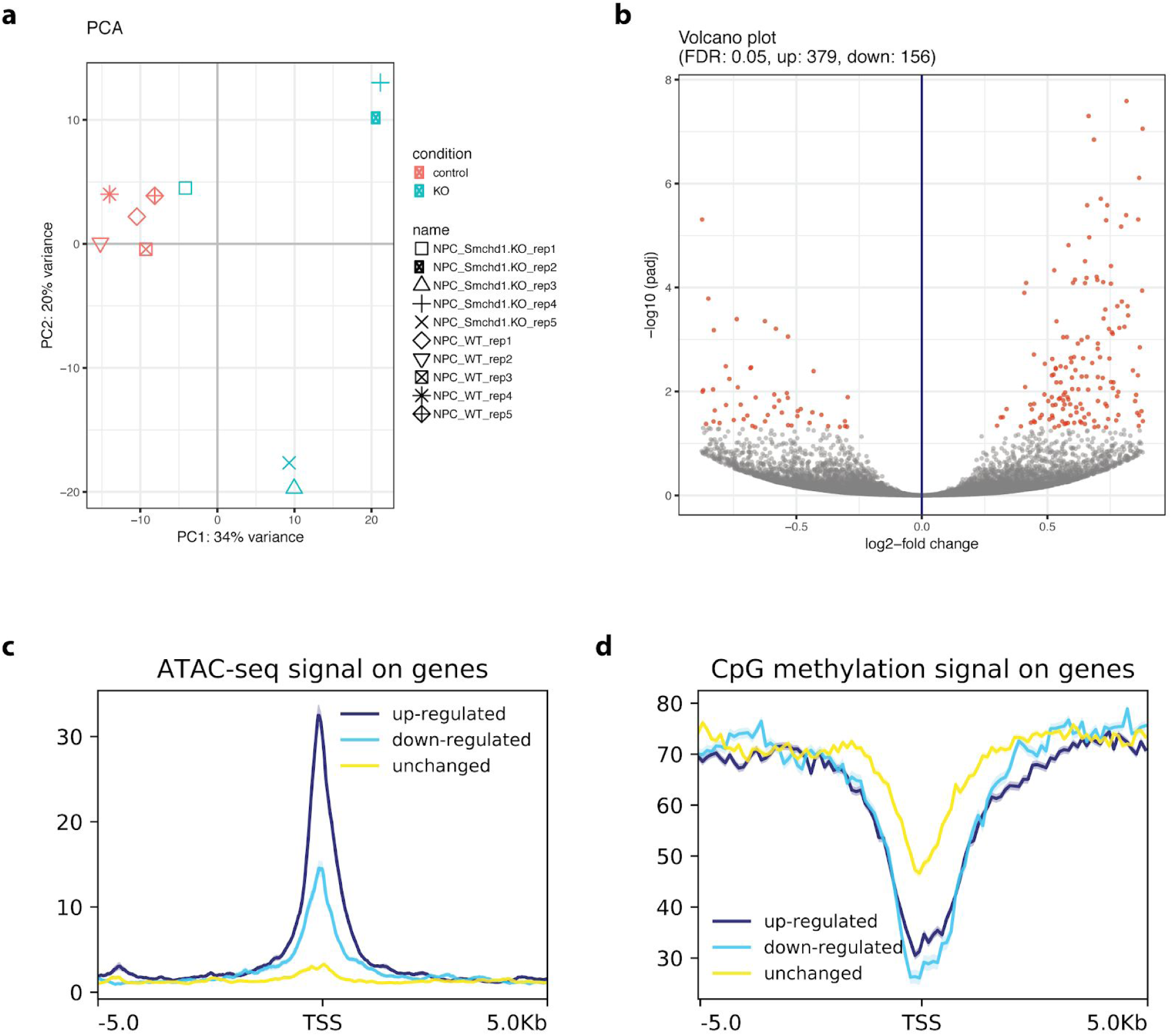
Analysis of de-repressed genes upon Schmd1 knock-out. A. PCA output from snakepipes suggested that one knock-out sample (replicate1) behaves differently. This sample was later revealed to be the XO clone which lost it’s inactive X chromosome. The sample was removed for DESeq2 analysis and the workflow was re-run. **B.** Volcano plot for DESeq2 output from snakePipes (knock-out replicate 1 excluded), shows an increase in up-regulated genes, indicating de-repression upon knock-out. **C.** Wild-type ATAC-seq signal on UP, DOWN and unchanged (NONE) genes, gene lists were extracted from DESeq2 output of RNA-seq workflow and depth-normalized bigwigs from ATAC-seq workflow was used for plotting. **D.** Wild-type methylation level reported by the WGBS workflow on UP, DOWN and unchanged (NONE) genes. (TSS = Transcription Start Site)

